# Staphylococcal DNA repair is required for infection

**DOI:** 10.1101/2020.02.23.961458

**Authors:** Kam Pou Ha, Rebecca S. Clarke, Gyu-Lee Kim, Jane L. Brittan, Jessica E. Rowley, Despoina A. I. Mavridou, Dane Parker, Thomas B. Clarke, Angela H. Nobbs, Andrew M. Edwards

## Abstract

The repair of DNA damage is essential for bacterial viability and contributes to adaptation via increased rates of mutation and recombination. However, the mechanisms by which DNA is damaged and repaired during infection are poorly understood. Using a panel of transposon mutants, we identified the *rexBA* operon as important for the survival of *Staphylococcus aureus* in whole human blood. Mutants lacking *rexB* were also attenuated for virulence in murine models of both systemic and skin infections. We then demonstrated that RexAB is a member of the AddAB family of helicase/nuclease complexes responsible for initiating the repair of DNA double strand breaks. Using a fluorescent reporter system, we were able to show that neutrophils cause staphylococcal DNA double strand breaks via the oxidative burst, which are repaired by RexAB, leading to induction of the mutagenic SOS response. We found that RexAB homologues in *Enterococcus faecalis* and *Streptococcus gordonii* also promoted survival of these pathogens in human blood, suggesting that DNA double strand break repair is required for Gram-positive bacteria to survive in host tissues. Together, these data demonstrate that DNA is a target of host immune cells, leading to double-strand breaks, and that repair of this damage by an AddAB-family enzyme enables the survival of Gram-positive pathogens during infection.

## Introduction

The integrity of bacterial DNA is essential for viability. However, the causes of DNA damage during infection and the mechanisms employed by bacteria to repair this damage are poorly understood, particularly in the case of Gram-positive pathogens such as *Staphylococcus aureus*. This pathogen is responsible for a raft of serious invasive infections including bacteraemia, infective endocarditis and osteomyelitis^1^. Despite a potent immune response, many infections become chronic or recurrent^2^, implying that either *S*. *aureus* does not experience DNA damage during infection or that it has efficient mechanisms for damage repair.

Entry of *S*. *aureus* into normally sterile tissues triggers the infiltration of neutrophils to control infection^3–5^. Neutrophils phagocytose *S*. *aureus*, and expose the bacterium to a cocktail of antimicrobial peptides and proteases^3,6^, reactive nitrogen species and the oxidative burst that consists of various reactive oxygen species (ROS)^7–12^. Whilst the contribution of each ROS to bactericidal activity is the subject of investigation, there is compelling evidence that the oxidative burst is crucial for control of *S*. *aureus* infection^9,12,13^. For example, individuals with chronic granulomatous disease (CGD) are particularly prone to staphylococcal infections because their neutrophils are defective for the oxidative burst^4,14^. In keeping with this, *S*. *aureus* survives better in mice defective for the oxidative burst relative to wild-type animals, whilst the treatment of human neutrophils with an inhibitor of the oxidative burst increased staphylococcal survival relative to untreated immune cells^5,9,15^. However, even when the oxidative burst is functional, there is evidence that some *S*. *aureus* cells can survive in neutrophils, which contributes to the progression of infection^16,17^.

Despite the importance of the oxidative burst in combatting staphylococcal infection, relatively little is known about how it kills the pathogen. Studies with single oxidants such as H_2_O_2_ indicate that the molecular targets of the oxidative burst are broad and include proteins, lipids and DNA^18^. To survive this damage, *S*. *aureus* employs several stress response regulators and repair systems. For example, previous work has shown that DNA damage caused by *S*. *aureus* exposure to H_2_O_2_ leads to the initiation of the DNA repair SOS response^19^, which facilitates the excision of damaged bases or repair of double-strand breaks (DSBs) by homologous recombination^20^. However, H_2_O_2_ is a sub-optimal model for the oxidative burst because it is typically used at concentrations that exceed those generated by the oxidative burst^21^. Therefore, it is unclear whether staphylococcal DNA is damaged by neutrophil-generated ROS, what the nature of this damage is, how it is repaired by *S*. *aureus*, and the impact of this damage on infection.

To address this gap in our knowledge, we examined mutants defective for DNA repair and found that a member of the AddAB helicase/nuclease family of enzymes was required for staphylococcal survival in blood and murine models of systemic and skin infections. We also demonstrated that this complex is required for the repair of DNA double strand breaks caused by the oxidative burst of neutrophils, which leads to the induction of the mutagenic SOS response. Similar complexes were required for the survival of infective endocarditis pathogens *Streptococcus gordonii* and *Enterococcus faecalis* in human blood, demonstrating that DNA damage repair is an important mechanism by which Gram-positive pathogens withstand host defences.

## Results

### RexAB is required for staphylococcal survival in host tissues

To determine whether DNA damage occurs under infection conditions, we assembled a panel of *S*. *aureus* mutants from the ordered NARSA transposon library^22^. Each mutant was defective for a different protein associated with DNA repair, which would enable the nature of any damage to be identified. We then assessed the contribution of each repair protein to bacterial survival in the host by measuring CFU counts of mutants in an *ex vivo* whole human blood model of infection. Previous characterisation of this model system by our group and others has shown that *S*. *aureus* is rapidly phagocytosed by neutrophils and exposed to the oxidative burst and other killing mechanisms^23–25^.

Wild type *S*. *aureus* JE2 survived relatively well in human blood, with >60% of the inoculum viable after 2 h (Supplementary Fig. S1). By contrast, survival of mutants defective for *rexA* or *rexB* was <5% of the inoculum (Supplementary Fig. S1). Since the transposon insertion in *rexB* contains a terminator sequence that prevents transcription of *rexA*, the second gene in the operon, *rexB* mutants are defective for both *rexB* and *rexA*, and are effectively *rexBA* mutants^22^. Therefore, we confirmed the importance of RexAB for the survival of two distinct strains (JE2 and SH1000) in blood by complementation of the *rexB* mutants with a plasmid containing the *rexBA* operon (p*rexBA*), which restored bacterial survival to wild type levels (Fig. 1A,B). By contrast, the survival of *rexB* mutants transformed with the vector alone (pEmpty) was not changed from that of the mutant (Fig. 1A,B).

**Figure 1.**
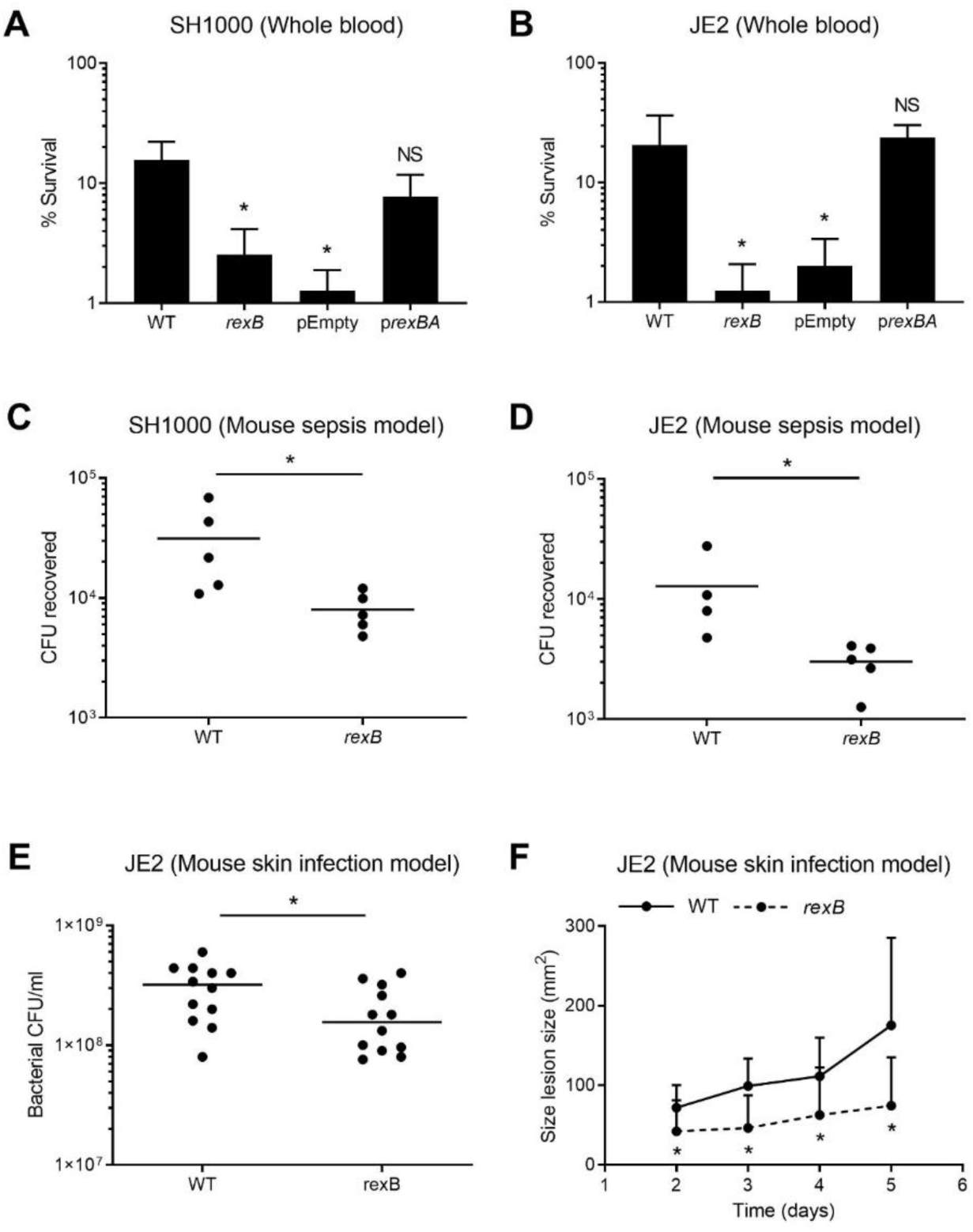
Survival of the *rexB* mutant is significantly reduced in human blood and *in vivo* murine infection models. **(A-B)** Survival of *S*. *aureus* wild type (WT), *rexB* mutant, empty vector (pEmpty) and complemented mutant (p*rexBA*) in SH1000 (A) and JE2 (B) backgrounds after 6 h incubation in whole human blood. Empty vectors and complemented mutants were supplemented with 100 ng/μl AHT to induce *rexBA* expression (n ≥ 3). **(C-D)** Number of CFU recovered from the peritoneal cavities of mice at 6 h post infection (each circle represents a single mouse; n = 4-5). **(E)** CFU/ml recovered from mouse skin biopsies at 5 days post infection (each circle represents a single mouse; n = 12). **(F)** Size of skin lesions observed on mice up to 5 days post infection (n = 12). Data in **A** and **B** were analysed by one-way ANOVA with Tukey’s post-hoc test relative to WT (*, P < 0.05). Data in **C-F** were analysed by Mann-Whitney test (*, P < 0.05).

Having shown that RexAB contributed to staphylococcal tolerance of neutrophil-mediated killing in human blood, we then tested its role in staphylococcal survival *in vivo* using a murine model of systemic infection. Mice were infected via the peritoneal cavity, which results in the recruitment of neutrophils within 2 h, with wild type or *rexB* mutants of *S*. *aureus* SH1000 or JE2^26^. After 6 h, the mice were sacrificed and the peritoneal cavity washed with PBS to recover bacteria, which were quantified by CFU counts. This revealed that *rexB* mutants in both genetic backgrounds were significantly attenuated for survival *in vivo*, with ∼5-fold lower CFU counts than the respective wild type bacteria (p ≤0.05), confirming that RexAB contributes to staphylococcal resistance to host immune defences and the progression of systemic infection (Fig. 1C,D).

Because *S*. *aureus* causes many different types of infection and the associated immune responses might vary, we next assessed survival of wild type strain JE2 and *rexB* mutant in a murine skin infection model. Mice were infected via sub-cutaneous injection and infection allowed to progress for 5 days before CFU counts at inoculation sites were determined. This revealed that wild type bacteria were present at 2-3-fold higher levels than the *rexB* mutant (p ≤0.05) (Fig. 1E). We also measured the size of the skin lesions generated by injected *S*. *aureus*. Wild type JE2 caused a lesion that progressively increased in size over time (Fig. 1F). By contrast, the lesion caused by the *rexB* mutant did not increase after day 2 and was significantly smaller than that caused by the wild type for days 2, 3, 4 and 5 (Fig. 1F; Supplementary Fig. S2). This indicated that RexAB was also required for skin infection progression.

To understand whether the reduced CFU counts of the *rexB* mutants relative to wild type in animal models were due to differences in growth rate or reduced virulence factor production, we measured bacterial growth, haemolysin production, staphyloxanthin levels and catalase across JE2 and SH1000 wild type and mutant strains. For both JE2 and SH1000, the *rexB* mutant did replicate at a slightly slower rate than wild type, but there were no significant differences in production of haemolysin, staphyloxanthin or catalase (Supplementary Fig. 3). Taken together, these findings demonstrate that RexAB significantly promotes staphylococcal survival in host tissues and is required for infection progression.

**Figure 2.**
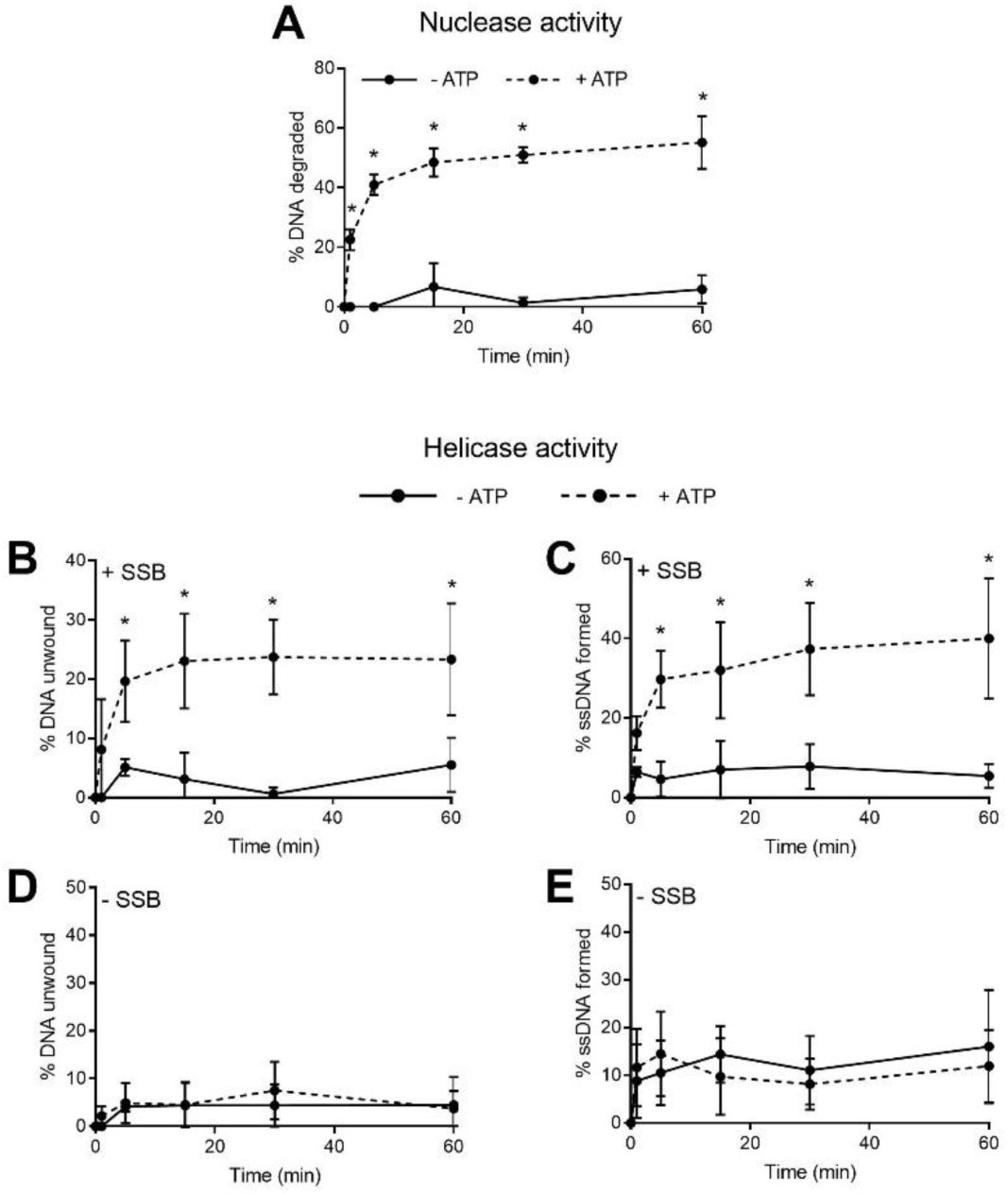
RexAB possesses ATP-dependent nuclease and helicase activity. **(A)** Nuclease activity of recombinant *S*. *aureus* RexAB protein in the presence (dotted line) or absence (solid line) of ATP (n = 4). **(B-E)** Helicase activity of recombinant *S*. *aureus* RexAB protein in the presence (dotted line) or absence (solid line) of ATP, by measuring unwinding of dsDNA and formation of ssDNA. Single-stranded DNA binding protein (SSB) was added to prevent reannealing of DNA (B-C) (n = 4). Equivalent assays were performed without SSB to confirm helicase activity (D-E) (n = 4). Data in **A-C** were analysed by two-way ANOVA with Sidak’s post-hoc test comparing without versus with ATP (*, P < 0.05). Error bars represent standard deviation of the mean.

**Figure 3.**
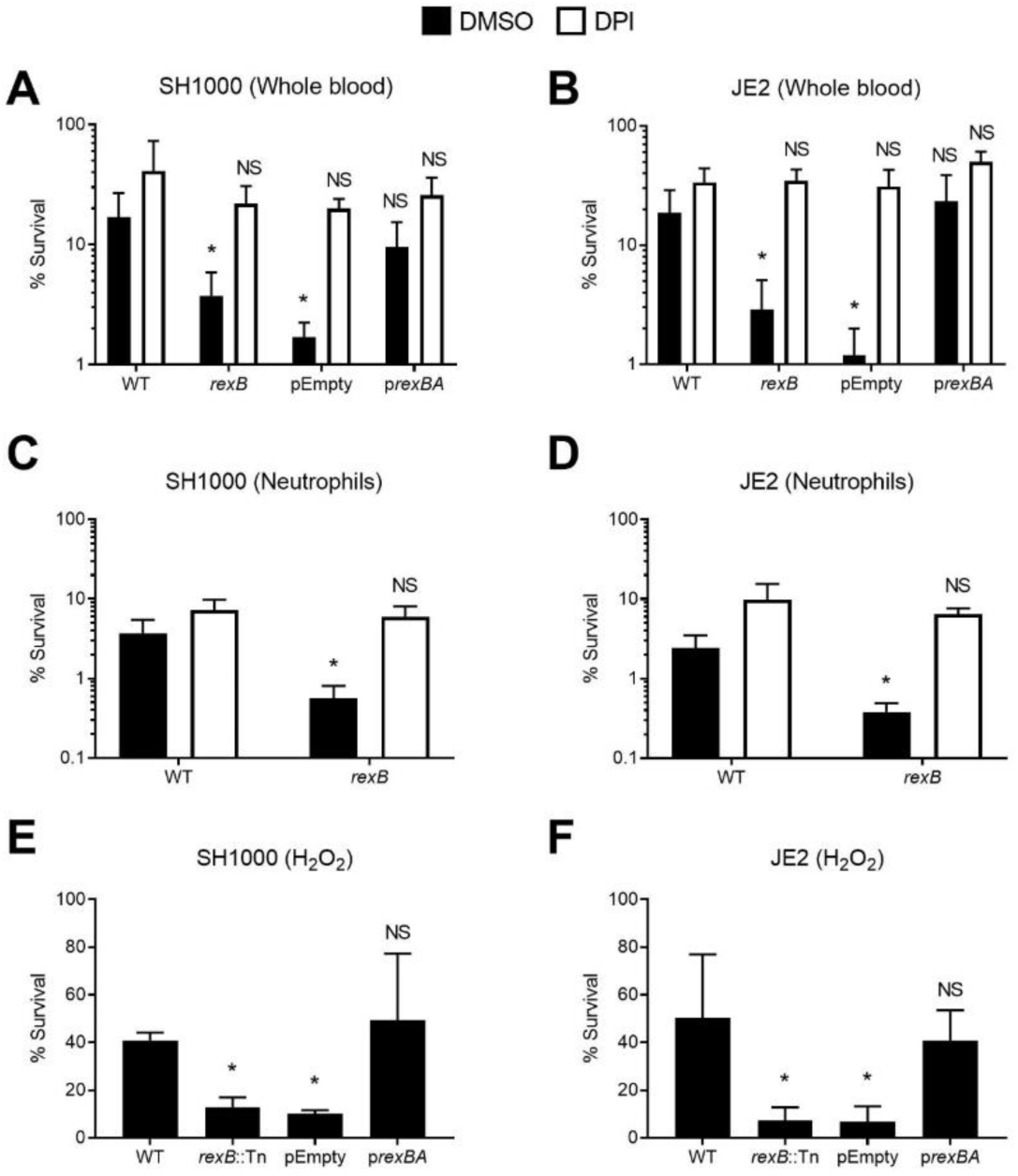
RexAB protects *S*. *aureus* from the oxidative burst of neutrophils and when exposed to H_2_O_2_. **(A-B)** Survival of *S*. *aureus* WT, *rexB* mutant, empty vector (pEmpty) and complemented mutant (p*rexBA*) in SH1000 (A) and JE2 (B) backgrounds in whole human blood after 6 h of incubation, in the presence of the oxidative burst inhibitor DPI (grey) or an identical volume of the DMSO solvent alone (black). Empty vectors and complemented mutants were supplemented with 100 ng/μl AHT to control *rexBA* expression (n ≥ 3). **(C-D)** Survival of *S*. *aureus* WT and *rexB* mutant when incubated with purified human neutrophils for 3 h (n = 4). **(E-F)** Survival of *S*. *aureus* WT, *rexB* mutant, empty vector (pEmpty) and complemented mutant (p*rexBA*) after 1 h incubation with 10 mM of H_2_O_2_. Empty vectors and complemented mutants were supplemented with 100 ng/μl AHT to induce *rexBA* expression (n = 3). Data in **A** and **B** were analysed by one-way ANOVA with Tukey’s post-hoc test relative to WT (*, P < 0.05). Data in **C** and **D** were analysed by Student’s t-test relative to WT (*, P < 0.05). Error bars represent standard deviation of the mean. Data in **E** and **F** were analysed by one-way ANOVA with Dunnett’s post-hoc test relative to WT (*, P < 0.05).

### RexAB is a member of the AddAB family of ATP-dependent helicase/nucleases

The *rexA* and *rexB* genes form a two gene operon (*rexBA*) on the staphylococcal chromosome, and are proposed to encode an AddAB helicase/nuclease enzyme on the basis of sequence homology^27–30^. However, this had not been demonstrated experimentally. Our initial *in silico* structural analysis of the predicted *rexBA* gene products supported predictions that this operon encodes an AddAB-type ATP-dependent helicase/nuclease that contributes to the processing and repair of DNA double strand breaks (Supplementary Fig. S4). AddAB enzymes process DSBs to generate a 3’ single-stranded DNA overhang that is necessary for RecA-mediated homologous recombination^31^. In support of the structural predictions, phenotypic testing of *rexBA* mutants showed that they were >8-fold more susceptible than wild type strains to the DNA-damaging antibiotics ciprofloxacin and mitomycin C, both of which cause DNA DSBs^32,33^ (Supplementary Fig. S5).

To confirm the ATP-dependent helicase/nuclease activity of the *S*. *aureus* RexAB complex, recombinant RexAB protein was generated and the helicase and nuclease activities were measured over 1 h. Nuclease activity assays were performed under conditions of high free Mg^2+^, which has been previously shown to activate nuclease activity in AddAB enzymes^34^. We found that DNA was degraded over time by the recombinant complex in the presence of ATP, whereas this degradation was minimal in its absence, demonstrating that RexAB has ATP-dependent nuclease activity (Fig. 2A).

We measured helicase activity at conditions of low free Mg^2+^ and observed loss of dsDNA, concomitant with increased ssDNA formation over time (Fig. 2B,C). These experiments were repeated in the absence of single-stranded DNA binding protein (SSB), which is required to prevent reannealing of DNA. In the absence of SSB, no DNA unwinding or ssDNA formation were observed, demonstrating that RexAB has ATP-dependent helicase activity (Fig. 2D,E). Combined, these results confirm that RexAB is a functional member of the AddAB family of DNA repair complexes. In turn, this demonstrates that DNA DSBs occur during staphylococcal infection and must be repaired for bacterial survival.

### RexAB enables staphylococcal tolerance of the oxidative burst

Having confirmed that DNA DSBs occur in *S*. *aureus* during infection, we wanted to determine whether this was due to the oxidative burst of neutrophils. Therefore, we incubated wild type and *rexB* mutants in whole human blood in the presence of diphenyleneiodonium chloride (DPI) to block the NADPH oxidase generated oxidative burst, or DMSO alone as a solvent control.

As already described in Fig. 1A,B, the survival of *rexB* mutants in whole human blood was significantly reduced relative to wild-type bacteria (Fig. 3A,B). However, the presence of DPI promoted the survival of *rexB* mutants to wild type levels, indicating that the survival deficit observed for bacteria lacking RexAB was due to increased sensitivity to the oxidative burst (Fig. 3A,B). To ensure that the killing of *rexB* mutants in blood was due to neutrophils, *S*. *aureus* strains were incubated with purified neutrophils and survival measured via CFU counts. Similar to whole blood, *rexB* mutants were more susceptible to neutrophil-mediated killing than wild type, but the presence of DPI restored survival of the *rexB* mutants to wild type levels (Fig. 3C,D). These data strongly indicated that RexAB contributed to staphylococcal survival of the oxidative burst by repairing DNA damaged by the oxidative burst.

To confirm that *rexB* mutants were more susceptible to oxidative damage, we measured survival of *S*. *aureus* strains in H_2_O_2_, which is one of the ROS produced in the oxidative burst^4^. As observed for whole human blood and purified neutrophils, *rexB* mutants were more susceptible to H_2_O_2_ than wild type or complemented strains.

Together, these data demonstrate that neutrophils cause DNA DSBs in *S*. *aureus* via the oxidative burst. This damage must be repaired by RexAB to enable staphylococcal survival in the host.

### RexAB is required for induction of the SOS response during exposure to the oxidative burst

Processing of DNA DSBs by AddAB proteins leads to the generation of a 3’ overhang. This results in the formation of a RecA filament, which triggers the SOS response, a multi-component DNA repair mechanism that mediates repair of the DNA DSB^20,35^. However, induction of the SOS response also leads to a transient increase in the mutation rate, which promotes the emergence of mutants with resistance to antibiotics or host-adapted phenotypes such as small colony variants^36,37^.

Therefore, we tested whether the neutrophil oxidative burst triggers the SOS response and whether this was dependent upon RexAB. To do this we used a P*recA-gfp* reporter construct and validated it by showing dose-dependent activity with mitomycin C, a well-established trigger of the SOS response^38^ (Supplementary Fig. S6). We then incubated TRITC-labelled *S*. *aureus* JE2 wild type and *rexB* mutants containing the reporter strain with neutrophils for 30 min and used flow cytometry to measure phagocytosis and reporter activity (GFP fluorescence) (Supplementary Fig. S7). As reported previously, >95% of *S*. *aureus* cells were phagocytosed by neutrophils within 30 min (Fig. 4A)^24^. Also, by 30 min, there was an increase in the GFP signal from wild type *S*. *aureus* relative to the start of the assay (Fig. 4B). By contrast, there was no increase in GFP signal from the *rexB* mutant (Fig. 4B), indicating that neutrophils trigger the SOS response in *S*. *aureus* via DNA processing by RexAB.

**Figure 4.**
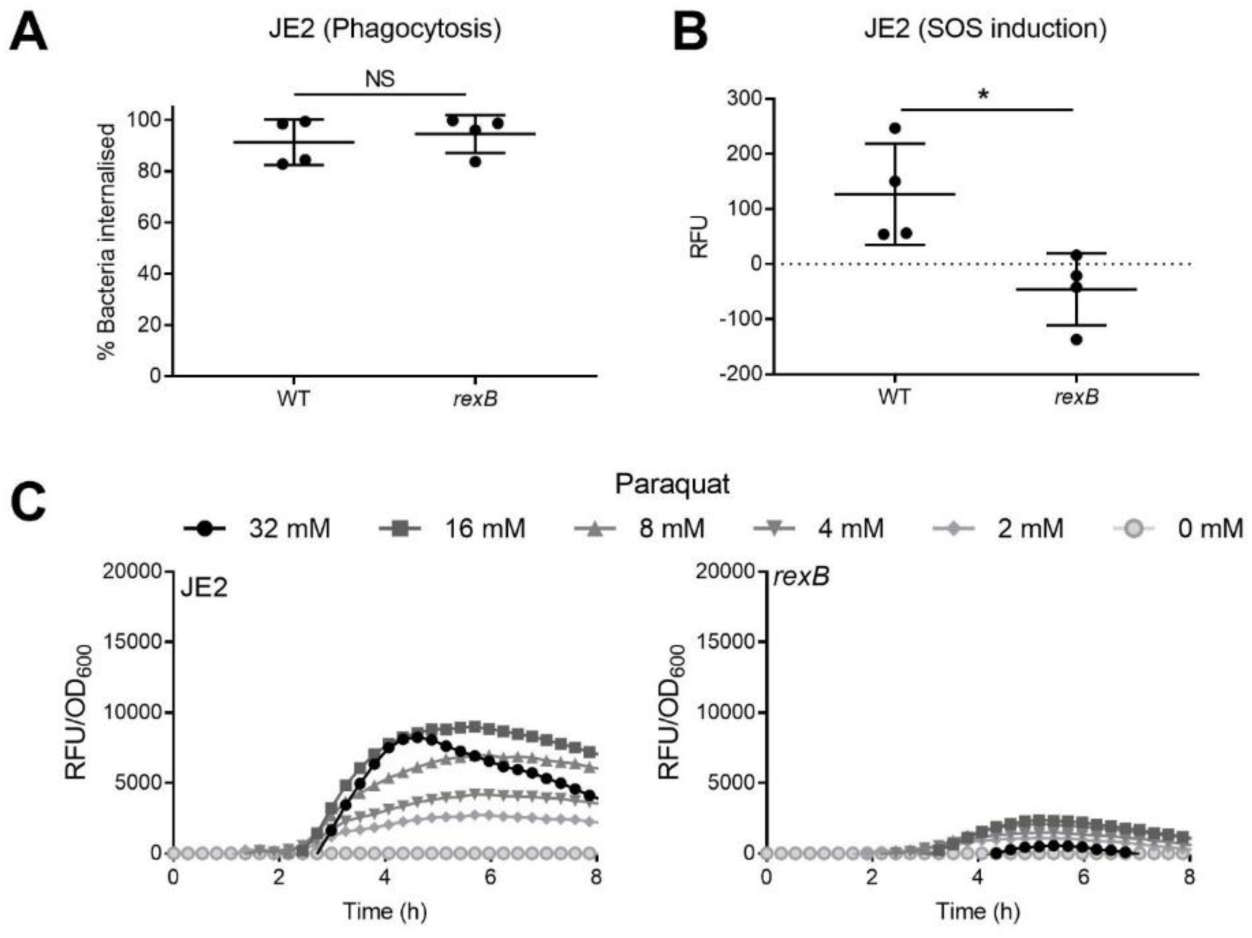
RexAB is required for induction of the SOS response during exposure to the oxidative burst of neutrophils and when exposed to the ROS generator paraquat. **(A)** Phagocytosis of JE2 wild type and *rexB* mutant *recA-gfp* reporter strains by neutrophils and **(B)** resulting induction of the SOS response measured by GFP expression (RFU) after 30 min of exposure (n = 4). **(C)** Induction of SOS response of JE2 wild type and *rexB* mutant as measured by GFP expression upon exposure to sub-lethal concentrations of paraquat. GFP fluorescence was normalised by OD_600_ to determine induction of SOS relative to cell density (n = 4). OD_600_ measurements alone are shown in Fig. S5. Data in **A** and **B** were analysed by Mann-Whitney test (*, P < 0.05). Where shown, error bars represent standard deviation of the mean. Error bars were omitted in **C** for clarity.

To further explore the requirement for RexAB for the induction of the SOS response during exposure to oxidative stress, wild type and *rexB* mutant bacteria were incubated with various sub-inhibitory doses of paraquat, which results in the generation of superoxide. For wild type bacteria, there was a clear dose-dependent increase in GFP-mediated fluorescence, indicative of SOS induction (Fig. 4C). For the *rexB* mutant, whilst there also appeared to be a dose-dependent induction of the SOS response, it was at considerably lower levels than that seen for the wild type (Fig. 4C). Therefore, the induction of SOS in response to oxidative stress is almost entirely dependent upon RexAB-mediated processing of DNA DSBs.

### RexAB is required for survival of streptococci and enterococci in human blood

Since homologues of RexAB are present in most Gram-positive bacteria^29–31^, we next tested whether this repair complex contributes to the survival of other bacteria exposed to neutrophils. Like *S*. *aureus, Enterococcus faecalis* and *Streptococcus gordonii* are frequent causes of infective endocarditis, which brings these species into close contact with neutrophils^39,40^. Therefore, *rexBA* was deleted in representative strains of each species and their sensitivity to the microbicidal activity of neutrophils determined using the *ex vivo* whole human blood model.

Wild type *E*. *faecalis* survived at high levels in human blood with ∼100% of the inoculum remaining viable during the full 6 h duration of the assay, but the loss of *rexBA* reduced enterococcal survival by ∼50% (Fig. 5A). This indicated that *E*. *faecalis* suffers DNA damage whilst in blood, but it can be tolerated via DNA repair (Fig. 5A). However, in contrast to *S*. *aureus*, DNA damage in *E*. *faecalis* was not due to the oxidative burst since survival of both wild type and *rexB* mutant in blood was unaffected by the presence of DPI (Fig. 5A).

**Figure 5.**
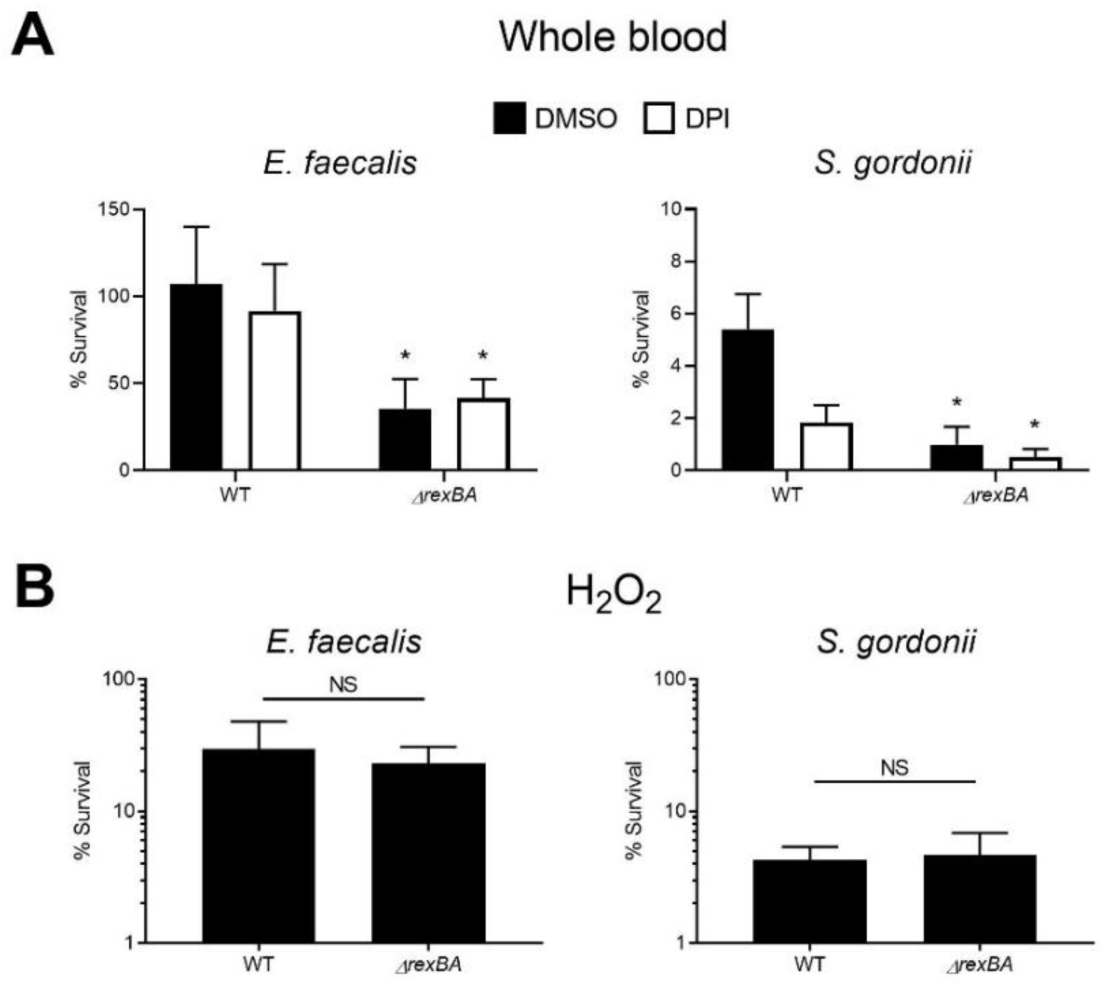
RexAB protects *Enterococcus faecalis* and *Streptococcus gordonii* from host defences in blood. **(A)** Survival of *E*. *faecalis* and *S*. *gordonii* wild type and *ΔrexBA* mutant in whole human blood after 6 h in the presence of the oxidative burst inhibitor DPI (white) or an identical volume of the DMSO solvent alone (black) (n = 3). **(B)** Survival of *E*. *faecalis* and *S*. *gordonii* wild type and Δ*rexBA* mutant exposed to H_2_O_2_ for 1 h (n = 3). Data in **A** and **B** were analysed by were analysed by Student’s t-test comparing *ΔrexBA* mutant versus WT (*, P < 0.05). Error bars represent standard deviation of the mean.

*S*. *gordonii* was much more susceptible to host defences in blood than *E*. *faecalis*, with ∼5% wild type bacteria remaining viable after 6 h (Fig. 5A). However, survival of the *rexBA* mutant was still reduced relative to the wild type, with ∼1% streptococci surviving after 6 h, indicating that DNA repair also contributes to the survival of *S*. *gordonii* in blood (Fig. 5A). As for *E*. *faecalis*, the presence of DPI did not increase the survival of *S*. *gordonii* in blood, indicating that DNA damage was not due to the oxidative burst in either bacteria. In support of these findings, the *rexBA* mutants of both *S*. *gordonii* and *E*. *faecalis* were no more susceptible to H_2_O_2_ than wild type bacteria (Fig. 5B).

Therefore, as for *S*. *aureus*, neutrophils damage DNA of both *S*. *gordonii* and *E*. *faecalis*, the repair of which by RexAB promotes bacterial survival. However, in contrast to *S*. *aureus*, the neutrophil oxidative burst does not appear to contribute to DNA damage in these bacteria.

## Discussion

Neutrophils are an essential host defence against *S*. *aureus* and many other bacterial pathogens. However, our understanding of the mechanisms by which these immune cells kill staphylococci is limited. The data presented here demonstrate that neutrophils cause DNA damage in *S*. *aureus* via the oxidative burst, the repair of which requires the RexAB complex and leads to induction of the SOS response. RexAB is a member of the AddAB helicase/nuclease family of enzymes involved in the processing of DNA DSBs for repair via homologous recombination. Thus, the oxidative burst of neutrophils causes DNA DSBs in *S*. *aureus*, which are lethal if not repaired. This provides new information on both the target of the oxidative burst, and the mechanisms by which *S*. *aureus* repairs and survives this damage. The importance of RexAB for staphylococcal survival during infection was demonstrated in murine models of both systemic and skin infection, as well as an *ex vivo* whole human blood model of bacteraemia. Since *S*. *aureus* is a frequent cause of both superficial and systemic infections^1^, these findings confirm the importance of DNA repair for staphylococcal survival in relevant host tissues.

Previous work has demonstrated the requirement of DNA DSB repair for the survival of Gram-negative pathogens *in vivo*. For example, AddAB was shown to be required for the infection of chickens and mice by *Campylobacter jejuni* and *Helicobacter pylori* respectively^41,42^. Furthermore, the virulence of *Salmonella enterica* in a murine model of bacteraemia was dependent upon the RecBCD DNA DSB repair complex^43^. However, the importance of such systems for Gram-positive bacteria was unclear.

DNA damage occurred within 30 min of phagocytosis of *S*. *aureus* by neutrophils, which corresponds to the time at which ROS are maximally generated in these immune cells^44,45^. The finding that the oxidative burst leads to DNA DSBs in *S*. *aureus* is in keeping with previous reports that ROS damage the DNA of Gram-negative pathogens such as *Escherichia coli, Salmonella enterica* and *Coxiella burnetii*, with the survival of bacteria phagocytosed by macrophages dependent upon RecBCD^43,46,47^. Therefore, DNA is a common target of the oxidative burst of phagocytic immune cells for several different human pathogens.

The RexAB system was also demonstrated to be important for survival of the infective endocarditis pathogens *E*. *faecalis* and *S*. *gordonii*. However, whilst neutrophils in blood caused DNA DSBs in these pathogens, this did not appear to be due to the oxidative burst, because the inhibition of NADPH oxidase with DPI had no effect on the survival of the wild type or *rexBA* mutants. Neutrophils employ several different antibacterial elements to kill invading pathogens, including reactive oxygen and nitrogen species, proteases and antimicrobial peptides. Several studies have indicated that bacteria vary in their susceptibility to each of these microbicides. In agreement with our findings, Standish *et al*. showed that *S*. *aureus* but not *Streptococcus pneumoniae* was killed by the oxidative burst^48^. However, whilst *S*. *aureus* can grow in the presence of nitric oxide, the replication of several other pathogens including *Pseudomonas aeruginosa* and *Streptococcus pyogenes* is inhibited^49^. Since reactive nitrogen species can cause DNA damage, this may provide an explanation for the finding that mutants of *E*. *faecalis* and *S*. *gordonii* lacking RexAB are more susceptible to killing by neutrophils in which the oxidative burst is blocked.

In addition to providing protection from oxidative damage caused by neutrophils, we have previously shown that RexAB provides tolerance to the combination antibiotic co-trimoxazole^38^. That work showed that DNA damage was partly due to endogenous oxidative stress that occurred during exposure to the combination antibiotic, in addition to direct damage to DNA caused by thymidine limitation^38^. The data presented here show that loss of RexAB also sensitises *S*. *aureus* to ciprofloxacin, even though the JE2 strain is resistant. In addition, there is growing evidence that multiple classes of antibiotics cause endogenous ROS production in *S*. *aureus*, suggesting that RexAB may provide an important defence against damage caused by both host defences and multiple classes of antibiotics, the two key threats to staphylococcal survival in the host. Further studies are needed to test this, but the identification of RexAB as important for staphylococcal survival during exposure to both neutrophils and at least two antibiotics makes this complex a potential target for novel therapeutics, particularly as the lack of RexAB homologues in eukaryotes reduces the likelihood of host toxicity^50^.

Inhibitors of AddAB and RecBCD have been previously reported in the literature, but problems include limited *in vivo* stability, poor oral bioavailability and a sub-optimal mechanism of action^51–54^. However, Amundsen *et al*. (2012) identified several small-molecule inhibitors of *Helicobacter pylori* AddAB and *E*. *coli* RecBCD, in particular ML328^55^, indicating that the development of stable, potent inhibitors is possible. More recently, a derivative of ML328 (IMP-1700) was found to be capable of potentiating antibiotic activity such that a resistant *S*. *aureus* strain was sensitised to ciprofloxacin^56^. However, the ability of IMP-1700 to inhibit AddAB/RecBCD activity under *in vivo* conditions remains to be determined. Further work in this area may lead to broad-spectrum therapeutics that promote bacterial susceptibility to both host defences and antibiotics as well as inhibiting induction of the mutagenic SOS response associated with the acquisition of drug resistance and host adaptation^37^.

In summary, staphylococcal, streptococcal and enterococcal DNA is damaged by the host immune system, leading to DNA DSBs that are lethal if not repaired by RexAB. These findings suggest that the RexAB complex is a potentially viable target for novel therapeutics, capable of sensitising Gram-positive pathogens to neutrophil-mediated killing and blocking the SOS response associated with the emergence of drug resistance.

## Materials and Methods

### Bacterial strains and culture conditions

The bacterial strains used in this study are listed in Table 1. *S*. *aureus* was cultured in Tryptic Soy Broth (TSB) to stationary phase (18 h) at 37 °C, with shaking (180 rpm). *S*. *gordonii* and *E*. *faecalis* were grown in Todd Hewitt Broth (THB) supplemented with 1% (w/v) yeast extract (THB-Y) at 37 °C, statically in 5% CO_2_. *E*. *coli* was grown in Lysogeny Broth (LB) or in Terrific Broth (TB) (1.2% (w/v) tryptone, 2.4% (w/v) yeast extract, 0.5% glycerol, 0.17 M KH_2_PO_4_, 0.72 M K_2_HPO_4_) for protein expression at 37 °C with shaking (180 rpm). Media were supplemented with antibiotics as required. When appropriate, bacteria were grown on Columbia Blood Agar (CBA) made with 5% defibrinated sheep blood.

**Table 1.** Bacterial strains used in this study.

| Strain | Description | Source |
| --- | --- | --- |
| ***Staphylococcus aureus*** |  |  |
| SH1000 | *rsbU*+ derivative of the laboratory strain 8325-4 | 57 |
| SH1000 *rexB*::Tn | SH1000 with a *bursa aurealis* transposon insertion in *rexB*, Ery<sup>r</sup> | 38 |
| SH1000 *rexB*::Tn *pitet* empty | SH1000 with a *bursa aurealis* transposon insertion in *rexB* with integrated *pitet* empty plasmid, Ery<sup>r</sup> | 38 |
| SH1000 *rexB*::Tn *pitet* *rexAB* | SH1000 with a *bursa aurealis* transposon insertion in *rexB* with integrated *pitet* with AHT-inducible *rexB*, Ery<sup>r</sup> | 38 |
| JE2 | A derivative of CA-MRSA USA300 LAC, cured of plasmids | 22 |
| JE2 *rexB*::Tn | JE2 with a *bursa aurealis* transposon insertion in *rexB*, Ery<sup>r</sup> | 22 |
| JE2 *rexB*::Tn *pitet* empty | JE2 with a *bursa aurealis* transposon insertion in *rexB* with integrated *pitet* empty plasmid, Ery<sup>r</sup> | 38 |
| JE2 *rexB*::Tn *pitet* *rexAB* | JE2 with a *bursa aurealis* transposon insertion in *rexB* with integrated *pitet* with AHT-inducible *rexB*, Ery<sup>r</sup> | 38 |
| JE2 pCN34 *PrecA-gfp* | JE2 containing pCN34 with *gfp* under the control of the *recA* promoter, Kan<sup>r</sup> | 38 |
| JE2 *rexB*::Tn pCN34 *PrecA-gfp* | JE2 *rexB*::Tn containing pCN34 with *gfp* under the control of the *recA* promoter, Kan<sup>r</sup> | 38 |
| ***Streptococcus gordonii*** |  |  |
| DL1 (Challis) | Wild type | 58 |
| DL1 Δ*rexBA* | DL1 with the *rexA* and *rexB* genes deleted | This study |
| ***Enterococcus faecalis*** |  |  |
| CK111 (pCF10-101) | Conjugative donor strain with pCF10-101 | 59 |
| OG1X | Gelatinase deficient | 60 |
| OG1X Δ*rexBA* | OG1X with the *rexA* and *rexB* genes deleted | This study |
| ***Escherichia coli*** |  |  |
| EC1000 | Cloning host that provides RepA in trans | 61 |
| SoluBL21 (DE3) | Derivative of BL21 (DE3) for the expression of challenging proteins | Genlantis |

### P*recA*-*gfp* fluorescent reporter assay

As detailed previously^38^, promoter-reporter gene constructs in the JE2 background was used to directly assess expression of *recA*. Antibiotic two-fold dilutions were made in flat-bottomed black-walled 96-well plates containing TSB and kanamycin (90 µg ml^-1^) and inoculated with 1/10 dilution of a stationary phase culture of the reporter strains. Plates were placed into an Infinite M200-PRO microplate reader (Tecan) where cultures were grown for 17 h at 37 °C (700 rpm), and both absorbance at 600 nm (OD_600_) and GFP relative fluorescence units (RFU) were measured every 30 min.

OD_600_ data and RFU data were normalised to no antibiotic controls. To account for differences in cell-density, RFU values were normalised by OD_600_ data at each time point.

### Neutrophil phagocytosis and measurement of DNA damage

Whole human blood (15 ml) was collected from individual healthy donors in EDTA-treated tubes (BD Biosciences) and layered over 20 ml of room-temperature Polymorph prep (Alere Limited) before centrifugation at 500 x g for 45-60 min (brake off, 30 °C) until a clear separation of red blood cells (RBCs), peripheral blood mononuclear cells (PBMCs) and polymophonuclear leukocytes (PMNs, or neutrophils) was seen. The PBMCs were discarded and the PMNs were transferred to a fresh centrifuge tube. Hanks’ Balanced Salt Solution (HBSS) was added to the PMNs to a total volume of 50 ml and cells were pelleted at 500 x g for 10 min (brake off, 30 °C). The cells were resuspended in 3 ml of HBSS, counted using a haemocytometer and adjusted to 5 × 10^6^ cells ml^-1^ in HBSS containing 10% human serum, 0.1 mM calcium and 0.1 mM magnesium. To the neutrophil suspension, 5 × 10^6^ cells ml^-1^ of bacteria (stationary or exponential-TRITC-stained) were added. The bacterial and neutrophil suspension was then incubated at 37 °C with tumbling. At each time point (0.5, 1, 2, 3 h) 100-150 μl was taken and resuspended in 4% PFA in PBS for a minimum of 1 h. Before analysis using flow cytometry, samples were washed and resuspended in PBS. Samples were analysed on the FACS Aria or LSRFortessa flow cytometer (BD Biosciences), and at least 100,000 events were captured, except for bacteria-only samples where at least 50,000 events were captured. Green (GFP bacteria) fluorescence was detected at 530 (30) nm, and TRITC labelling at 586 (16) nm. Full gating strategies are detailed in Figures S6 and S7.

### Determination of minimum inhibitory concentration (MIC)

Minimum inhibitory concentrations (MICs) were determined using a serial broth dilution protocol as described previously^62^. Bacteria were diluted to 1 × 10^5^ CFU ml^-1^ and incubated in flat-bottomed 96-well plates with a range of antibiotic concentrations for 17 h at 37 °C under static conditions (aerobic, anaerobic or 5% CO_2_). Media containing daptomycin was supplemented with 1.25 mM CaCl_2_. The MIC was defined as the lowest concentration at which no growth was observed.

### Whole blood and hydrogen peroxide survival assays

Bacteria were washed twice with PBS, adjusted to 10^6^ CFU ml^-1^ in HBSS and 10^4^ CFU (10 μl) used to inoculate 90 μl of freshly donated human blood (collected in EDTA-treated tubes; BD Biosciences) or freshly diluted H_2_O_2_ (10 mM in PBS) in 96 well plates. Ethical approval for drawing and using human blood was obtained from the Regional Ethics Committee and Imperial NHS Trust Tissue Bank (REC Wales approval no. 12/WA/0196, ICHTB HTA license no. 12275). In some assays blood was pre-treated for 10 min with diphenyleneiodonium (DPI; 50 µM) or an equivalent volume of DMSO as a solvent control. After 6 h incubation, bacterial survival was determined by CFU counts of blood-bacteria mixtures on CBA plates. For H_2_O_2_ assays, survival was measured after 1 h at 37 °C (static) in the dark Survival for both assays was calculated as a percentage of the number of bacteria in the starting inoculum.

### Neutrophil survival assay

Neutrophils were adjusted to 5 × 10^6^ cells ml^-1^ in HBSS containing 10% human serum, 0.1 mM calcium and 0.1 mM magnesium. Stationary phase bacterial cultures were washed in PBS and 1 × 10^6^ CFU was added to the neutrophil suspension (MOI 1:5) to a total volume of 1 ml. Neutrophils were treated for 10 min prior to the addition of bacteria with either diphenyleneiodonium (DPI; 50 µM) or an equivalent volume of DMSO (solvent control) as needed. The bacterial and neutrophil suspension was subsequently incubated at 37 °C with tumbling. At relevant time points (0.5, 1, 2, 3 h), 50 µl of the suspension was transferred to a 96-well plate and serially diluted ten-fold in PBS up to 10^−3^ dilution. All dilutions (including neat) were then plated onto CBA and incubated for 24 h at 37 °C before counting. Survival was calculated as a percentage of the number of bacteria in the starting inoculum.

### Murine systemic infection model

Animal work was conducted in accordance with the Animals (Scientific Procedures) Act 1986 outlined by the UK Home Office regulations. Work was approved by the UK Home Office after ethical approval by the Imperial College Animal Welfare and Ethical Review Body (AWERB). Six-to eight-week-old female C57BL/6 mice (Charles River) were infected via the intraperitoneal route with wild type or *rexB*::Tn mutant in JE2 and SH1000 backgrounds. Stationary phase bacterial cultures were washed twice with PBS and adjusted to 10^7^ CFU ml^- 1^. Subsequently, 400 μl (4 × 10^6^ CFU) of the washed bacterial suspensions were injected into the peritoneal cavity of each mouse (5 mice for each strain, 20 in total). After 6 h, the mice were humanely sacrificed by cervical dislocation and death confirmed by severing the femoral artery. The peritoneal cavity was washed with PBS to release the bacteria and CFU counts were determined by plating onto TSA. Sample size was determined prior to the experiment using power analysis based on *in vitro* data^63^. Tubes containing the bacterial suspensions were blinded before starting the experiment. Mice were randomly allocated to group cages and each group was randomly allocated to a treatment. Following Home Office regulations, any animals that displayed two or more of the following symptoms were humanely killed using a Schedule 1 method and excluded from the study: shivering, hunched posture, reduced movement, cyanosis, circling or difficulty breathing.

### Murine skin infection model

Six-to -eight-week-old mice were subcutaneously infected with 2 × 10^6^ CFU of exponential phase cultures of *S*. *aureus* as previously described^64^. Skin infection animal work was performed according to the Guidelines for the Care and Use of Laboratory Animals of the National Institutes of Health, the Animal Welfare Act, and U.S. Federal law. Protocol was approved by the Animal Care and Use Committee of Rutgers New Jersey Medical School.

### Construction of Δ*rexBA* mutants of *E*. *faecalis* and *S*. *gordonii*

A Δ*rexBA* mutant was generated in *S*. *gordonii* by in-frame allelic replacement with the erythromycin resistance determinant *ermAM*. Flanking regions directly upstream and downstream of the *rexBA* operon were amplified by PCR from *S*. *gordonii* DL1 genomic DNA with primer pairs Sg.rexAB.F1/Sg.rexAB.R1 and Sg.rexAB.F2/Sg.rexAB.R2 (Table 2), respectively, while the *ermAM* cassette was amplified from plasmid pVA838^65^ using primers ermAM.SgF/ermAM.SgR (Table 2). The resulting amplimers were then joined together via 20-bp overlapping regions by stitch PCR using primers Sg.rexAB.F1/Sg.rexAB.R2, and transformed into *S*. *gordonii*. Erythromycin-resistant transformants were confirmed by sequencing and the strain designated UB3018.

**Table 2.**
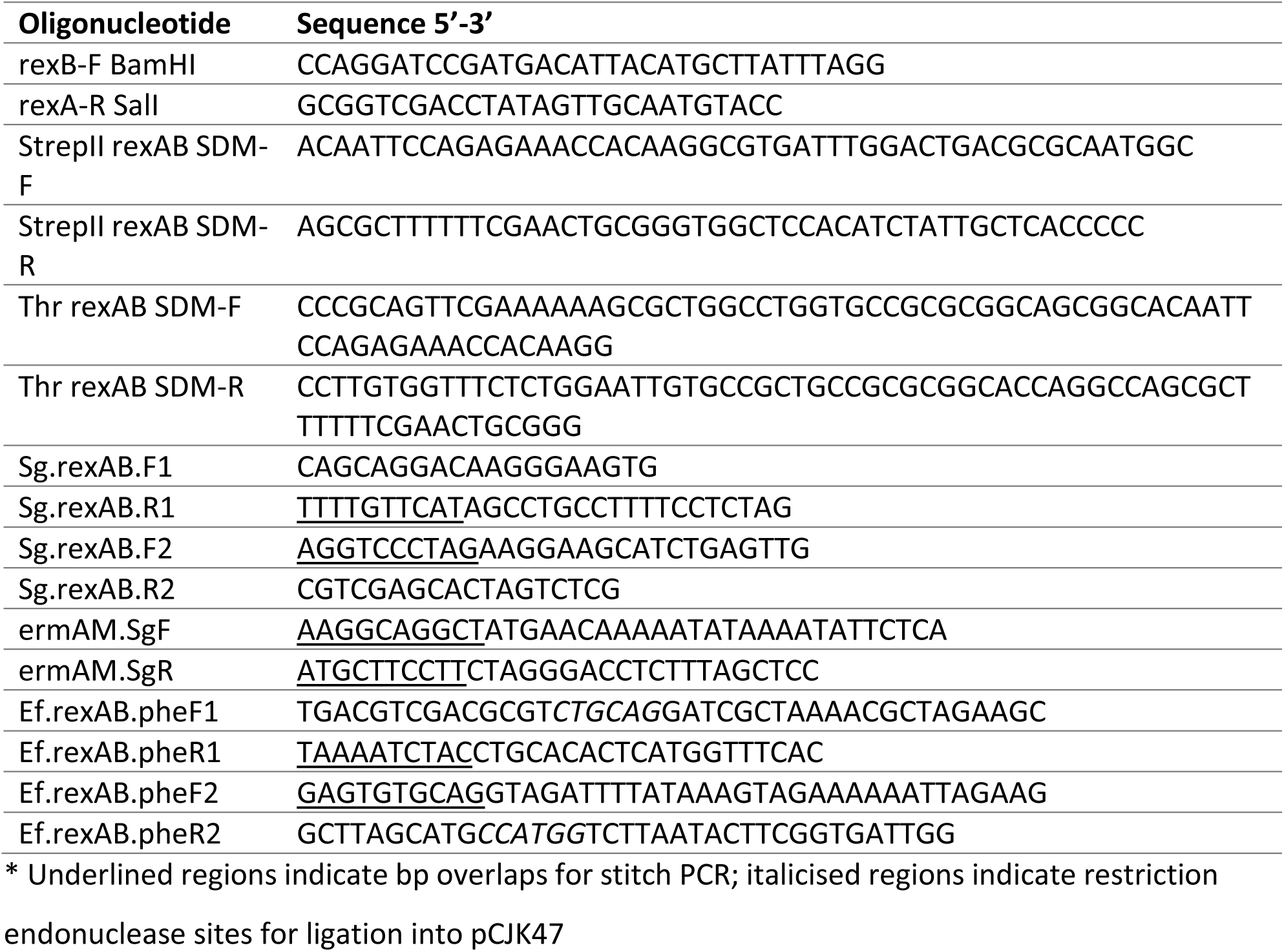
Primers used in this study.

A *rexBA* mutant was generated in *E*. *faecalis* by markerless exchange using a 2-step homologous recombination approach, as previously described^59^. In brief, flanking regions directly upstream and downstream of the *rexBA* operon were amplified by PCR from *E*. *faecalis* OG1X genomic DNA with primer pairs Ef.rexAB.pheF1/Ef.rexAB.pheR1 and Ef.rexAB.pheF2/Ef.rexAB.pheR2 (Table 2), respectively, and then joined together via 20-bp overlapping regions by stitch PCR using primers Ef.rexAB.pheF1/Ef.rexAB.pheR2. The resultant amplimer was cloned into donor plasmid pCJK47^59^ via unique restriction sites *Pst*I/*Nco*I to generate pCJK47-*rexAB*, propagated in *E*. *coli* EC1000 and then introduced into conjugative donor strain *E*. *faecalis* CK111 (pCF10-101) by electroporation. Plasmid pCJK47-*rexAB* was transferred to *E*. *faecalis* OG1X by conjugation. Transconjugants carrying the integrated plasmid were confirmed by colony PCR, before counterselection based on the P-*pheS*\* marker was used to identify secondary recombinants in which the integrated plasmid had been excised and lost, leaving the desired Δ*rexBA* allele. This was confirmed by sequencing and the strain designated UB2948.

### Construction of *S*. *aureus* RexAB expression vector

Cloning of the *rexA* and *rexB* genes from *S*. *aureus* was achieved by PCR from wild-type genomic JE2 DNA using the “rexB-F BamHI” and “rexA-R SalI” primers listed in Table 2, which allowed the amplification of the *rexBA* operon immediately flanked by suitable restriction endonuclease recognition sequences (BamHI and SalI). The *rexA* and *rexB* genes were inserted into the pET28b^+^ vector (Novagen) using standard cloning techniques and site-directed mutagenesis (SDM) was performed to insert a StrepII-tag and a thrombin site in front of the *rexA* gene. This enabled RexA and RexB proteins to be detected individually, via an N-terminal His_6_-tag for RexB (His_6_ from the pET28b^+^ vector) and an N-terminal StrepII-tag for RexA. PCR primers for SDM are listed in Table 2. DNA sequencing was used on the “pET28b^+^ *rexBA*” expression plasmid to confirm that the sequences of the entire *rexA* and *rexB* genes, tags and promoter regions were as expected.

### Expression and purification of recombinant *S*. *aureus* RexAB

Cells from single colonies of *E*. *coli* SoluBL21 (DE3), freshly transformed with pET28b^+^ *rexAB* expression plasmid coding for N-terminally StrepII-tagged RexA and N-terminally His_6_-tagged RexB, were used to inoculate an overnight starter culture in LB supplemented with 50 μg ml^- 1^ of kanamycin. The starter culture was diluted to an OD_600_ of 0.05 into 4 L of TB containing 50 μg ml^-1^ of kanamycin. Cells were grown at 37 °C with shaking at 180 rpm until an OD_600_ of 0.5 was reached prior to induction with 1 mM of isopropyl β-D-thiogalactoside (IPTG). Following induction, the temperature was reduced to 20 °C and cultures were further incubated for 20 h. Cells were harvested by centrifugation at >10,000 x *g* and 4 °C for 30 min and pellets were resuspended in 100 ml of 50 mM Tris, 150 mM NaCl (pH 7.5).

For protein purification, cells were disrupted by sonication and cell debris was cleared by centrifugation at 32,000 x g 4 °C and for 30 min. The resulting supernatant was added to 5 ml of Chelating Sepharose Fast Flow resin (GE Healthcare), which had been loaded with 0.1 M NiCl_2_ and equilibrated with 100 ml of 50 mM Tris, 150 mM NaCl, 20 mM imidazole (pH 7.5). The supernatant-resin mixture was left at 4 °C overnight with gentle stirring to optimize the binding of the His-tagged protein to the nickel-charged resin. Then, the mixture was washed 9x with 40 ml of 50 mM Tris, 150 mM NaCl, 20mM imidazole (pH 7.5) and 1x with 20 ml of 50 mM Tris, 150 mM NaCl, 70 mM imidazole (pH 7.5). His-tagged protein was eluted with 50 mM Tris, 150 mM NaCl, 150 mM imidazole (pH 7.5); each 10 ml fraction was tested with Bradford reagent (Bio-Rad) for protein content until no more protein could be detected. Fractions containing protein were pooled, buffer exchanged and concentrated using an Amicon 100 kDa cut-off concentrator (Merck Millipore); RexAB is >250 kDa in size. The concentrating device was centrifuged at 2000 x g (4 °C), and the protein solution was exchanged into 50 mM Tris, 150 mM NaCl (pH 7.5) by four serial concentration and re-dilution steps. The total protein concentration was quantified using the Pierce BCA protein assay kit (Thermo Fisher Scientific) following the manufacturer’s instructions, and presence of intact recombinant RexAB was confirmed via SDS-PAGE and Western blot analysis.

### Nuclease and helicase activity assays

The nuclease and helicase activity of RexAB was measured to confirm the AddAB-like activity in our recombinant RexAB protein. Staphylococcal DNA was amplified from the JE2 whole genome by colony PCR using the primer pairs “Chi control F” (5’-TCAGTGAATTAGATGATTCGC -3’) and “Chi control R” (5’-TTCATACGTATGAATGTTATTTGC-3’) where the amplicon lacked a Chi site region to be used as the DNA substrate in these assays.

Reactions were set up in either nuclease assay buffer (25 mM Tris-acetate pH 7.5, 2 mM Mg acetate, 1 mM DTT) or helicase assay buffer (25 mM Tris-acetate pH 7.5, 0.25 mM Mg acetate, 1 mM DTT), along with 5 ng µl^-1^ of DNA, 1 mM ATP or equivalent volume of nuclease-free water and either 20 nM or 50 nM of recombinant RexAB for the nuclease and helicase assay respectively. Additionally, for the helicase assay, 2 μM of SSB protein was added to each sample. The DMSO content was maintained at 1% (v/v) when compounds were being tested. Samples were incubated statically at 37 °C and at 0, 5, 15, 30, 60 and 120 min, 5 µl was removed and pipetted into 20 µl of STEB buffer (40 % (w/v) sucrose, 100 mM Tris-HCl, 10 mM EDTA, 0.5 mg/ml bromophenol blue, pH 8) to stop the reaction. 20 µl of chloroform/isoamyl alcohol (24:1) was added to each tube, vortexed for 10 sec and centrifuged for 2 min at 17,000 x g to remove the protein and any compounds used for inhibition. The aqueous (upper blue) phase was loaded onto a 1% (w/v) agarose gel prepared in TBE buffer and electrophoresis was carried out at 85 V for 1 h. The gels were subsequently stained with SYBR Safe DNA gel stain (Invitrogen) at 1/10,000 dilution in TBE buffer for 2 h with rocking and visualised using a Gel Doc EZ Imager (Bio-Rad). Band intensity was quantified using ImageJ software. For nuclease activity, values were normalised to no ATP controls at 0 h. For helicase activity, values were normalised to a ssDNA control that lacked the RexAB protein, which had been heated at 95 °C for 2 min to denature the dsDNA and allow the SSB protein to bind and stabilise the two ssDNA strands.

### Measurement of bacterial growth

To measure growth of *S*. *aureus*, bacterial cultures were first grown to stationary phase in TSB at 37 °C (180 rpm), then inoculated 1/50 for growth curves, or 1/10 for growth inhibition assays (supplementary) into a flat-bottomed 96-well plate (200 µl total volume) and placed into a POLARstar Omega plate reader (BMG Labtech). Bacteria were grown for 17 h at 37 °C (700 rpm), and absorbance at 600 nm was measured every 30 min.

### Haemolytic activity

The haemolytic activity of culture supernatants was determined as described previously^66^. Briefly, stationary phase *S*. *aureus* cultures were pelleted for 5 min at 17,000 x g, and 400 µl of the supernatant pipetted into microcentrifuge tubes. An equal volume of 2% defibrinated sheep blood in PBS was added and the mixture incubated statically for 1 h at 37 °C. Fresh TSB containing 2% defibrinated sheep blood was used as a negative control. After incubation, the samples were centrifuged for 5 min at 500 x g and 200 µl of supernatant was transferred into a flat-bottomed 96-well plate. The absorbance of released haemoglobin was measured at 540 nm using a POLARstar Omega plate reader (BMG Labtech). Percentage haemolysis was calculated relative to the wild type control.

### Staphyloxanthin extraction and quantification

To extract and quantify staphyloxanthin, *S*. *aureus* stationary phase cultures grown in TSB were harvested by centrifugation at 17,000 x g for 2 min. Staphyloxanthin was extracted by incubating the culture in methanol at 42 °C for 30 min. Cells were pelleted by centrifugation at 17,000 x g for 2 min, and 100 µl of supernatant was transferred into a flat-bottomed 96-well plate. The released staphyloxanthin was quantified by measuring the absorbance at 462 nm using a POLARstar Omega plate reader (BMG Labtech).

### Catalase activity

To determine the level of catalase activity, overnight cultures were washed three times in PBS and 10^7^ CFU were inoculated into 1 ml of 100 µM H_2_O_2_ (diluted in PBS). Samples were incubated at 37 °C protected from light for 15 min. 200 µl of the sample was centrifuged at 17,000 x g for 3 min, and 20 µl of the supernatant was added to a 96-well microtitre plate. PBS containing no bacteria was used as a negative control.

Catalase activity was determined indirectly by measuring the concentration of H_2_O_2_ over time, using the Pierce Quantitative Peroxide Assay kit (aqueous-compatible formulation) (Thermo Fisher Scientific). Following the manufacturer’s instructions, reagents A and B were mixed at a ratio of 1:100 and 200 µl was added to each sample in the 96-well plate. The plate was then incubated for 30 min at room temperature and the absorbance was measured at 595 nm using an iMark microplate reader (Bio-Rad). The H_2_O_2_ concentration was determined using a standard curve of known concentrations (up to 1 mM).

### Computational analyses

Multiple sequence alignments were generated using Clustal Omega via the EMBL-EBI web server^67^. Parameters were left in their default settings with the exception of alignment “order”, which was set to consider the input order. Protein structures were predicted using the Phyre2 protein fold recognition server^68^. Phyre2 structural models were viewed and manipulated using the PyMOL Molecular Graphics System Version 2.3 (Schrödinger).

### Statistical analyses

Data are represented as the mean or median from three or more independent experiments and analysed by Student’s t test (two tailed, unpaired, assuming equal variances), one-way ANOVA or two-way ANOVA corrected for multiple comparison, as described in the figure legends. For each experiment, “n” refers to the number of independent biological replicates. CFU counts from murine experiments are presented as the value obtained from each animal and significance was assessed using the Mann-Whitney test. P < 0.05 was considered significant between data points (GraphPad Prism 7 for Windows).

## Acknowledgements

A.M.E. and R.S.C. acknowledge funding from Shionogi & Co., Ltd. A.M.E. also acknowledges support from the National Institute for Health Research (NIHR) Imperial Biomedical Research Centre (BRC). K.P.H. is supported by a PhD scholarship funded by a Medical Research Council award to the Centre for Molecular Bacteriology and Infection (MR/J006874/1). D.P. is supported by the National Institutes of Health grants R01HL134870 and R21AI153646. D.A.I.M acknowledges funding from the Medical Research Council (Career Development Award MR/M009505/1). All authors acknowledge the provision of strains by the Network on Antimicrobial Resistance in Staphylococcus aureus (NARSA) Program: under NIAID/ NIH Contract No. HHSN272200700055C. The funders had no role in the study design, interpretation of the findings or the writing of the manuscript.

