## Supplementary information for "Staphylococcal DNA repair is required for infection"

Supplementary figures S1-S7

Supplementary references

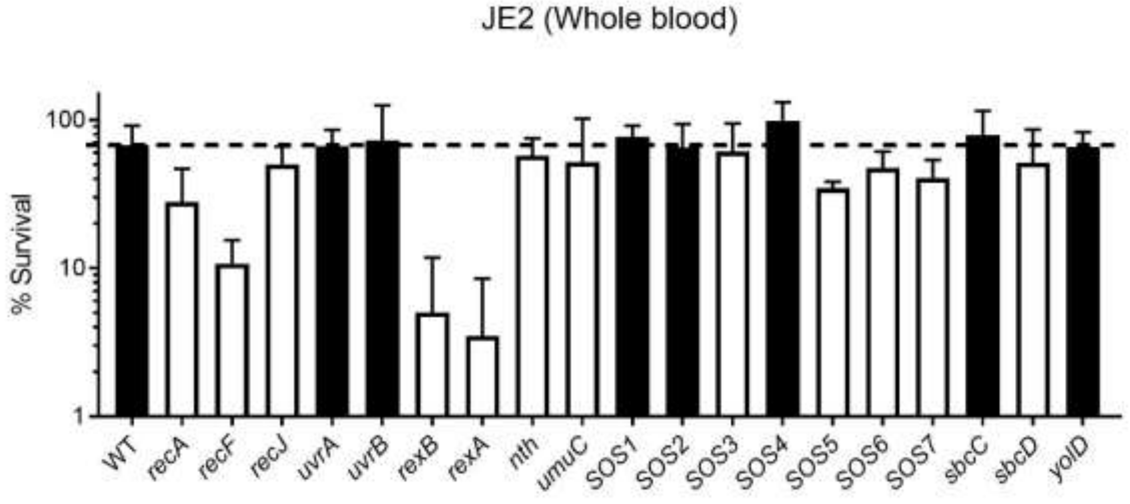


**Figure S1. Mutants lacking functional copies of *rexA* and *rexB* have the lowest survival of a range of DNA repair mutants in whole human blood.** Survival of *S. aureus* JE2 wild type and a number of DNA repair mutants from the Nebraska transposon mutant library in whole human blood after 2 h of incubation (n ≥ 4). SOS1-7 refers to genes of unknown function within the *S. aureus* SOS regulon. (SOS1: SAUSA300_0365/ SACOL0436; SOS2: SAUSA300_1238/ SACOL1330; SOS3: SAUSA300_1903/ SACOL1986; SOS4: SAUSA300_1915/SACOL1999; SOS5: SAUSA300_2129/SACOL2160; SOS6: SAUSA300_2130/SACOL2161; SOS7: SAUSA300_2131/SACOL2162). Mutants with lower percentage survival than the JE2 WT (represented by the dotted line) are shown in white. Error bars represent standard deviation of the mean.

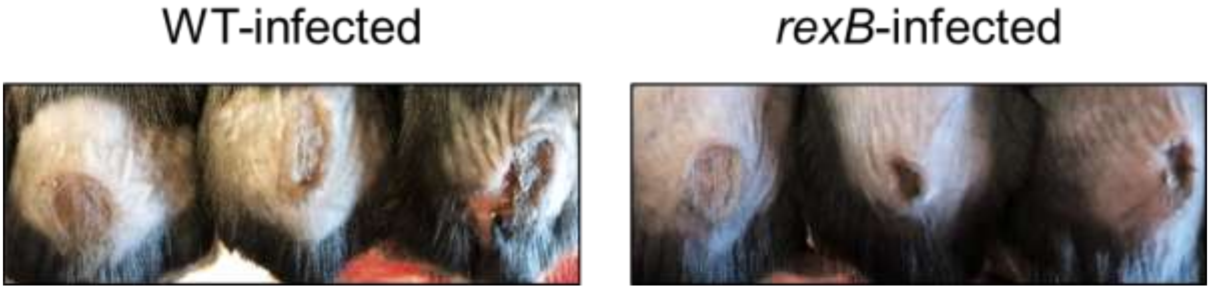


**Figure S2. Photographs of skin lesions caused by *S. aureus* wild type and the *rexB* mutant.** Representative images of skin lesions from WT- and *rexB*-infected mice at 5 days post infection.

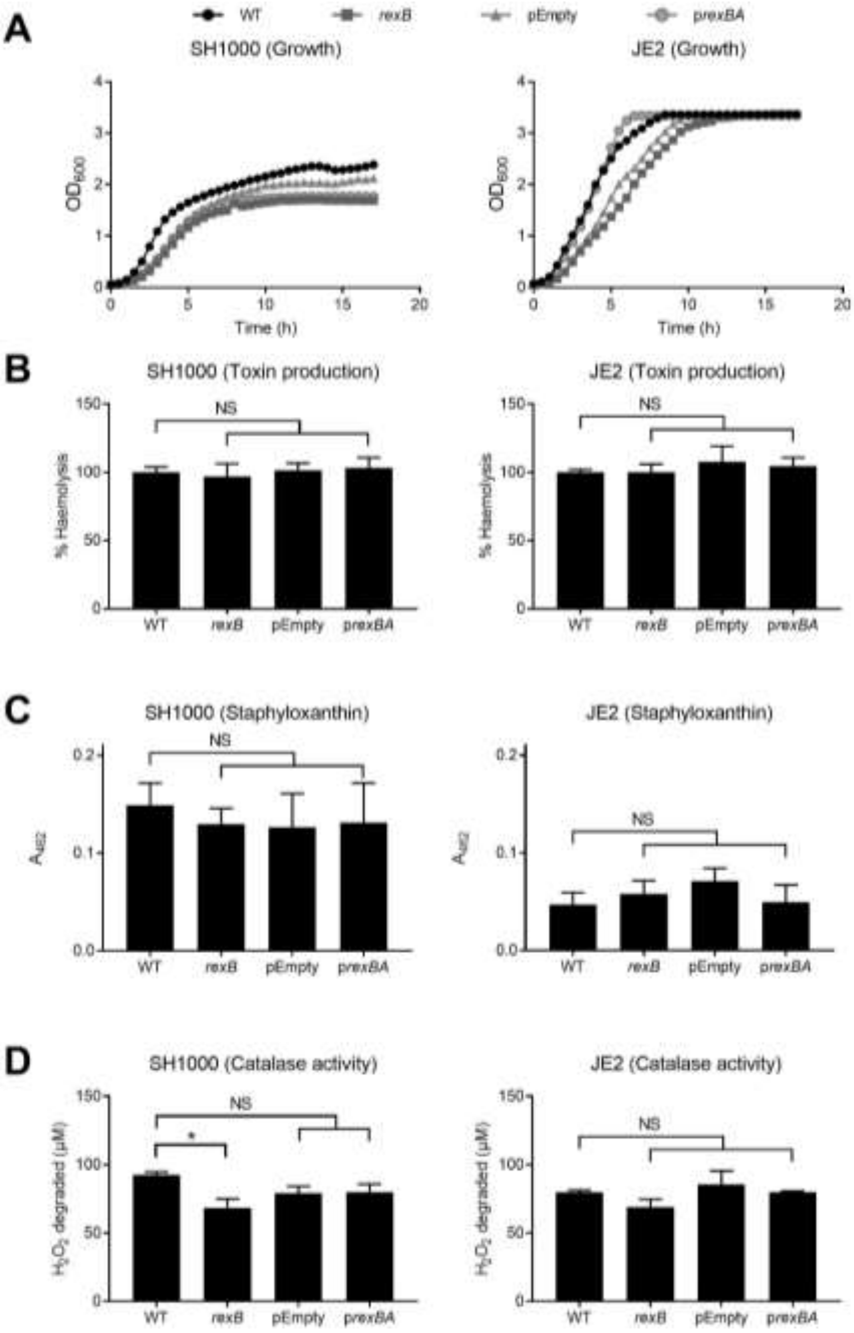


**Figure S3. Phenotypic properties of wild type and *rexB* mutant. (A)** Growth curves of WT, *rexB* mutant, empty vector (pEmpty) and complemented mutant (p*rexBA*) in TSB in SH1000 and JE2 backgrounds. TSB was supplemented with 100 ng/ml of AHT for pEmpty and p*rexBA* to control expression of *rexAB* (n = 3). **(B-D)** Graphs showing toxin production (B), staphyloxanthin production (C) and catalase activity (D) of WT, *rexB*, empty vector (pEmpty) and complemented mutant (p*rexBA*) in JE2 and SH1000 backgrounds. Cultures of pEmpty and p*rexBA* were grown in the presence of 100 ng/ml of AHT (n ≥ 3). Toxin production was determined by measuring haemolytic activity as an indicator for *agr* operon expression. Catalase activity measured the degradation of $H_2O_2$ after 15 min incubation with *S. aureus* cultures, which detected significance for SH1000 WT versus *rexB,* but this was not seen for SH1000 WT versus pEmpty or in JE2. Data were analysed by one-way ANOVA with Tukey's post-hoc test (*, $P < 0.05$). Where shown, error bars represent standard deviation of the mean. Error bars were omitted in **A-B** for clarity.

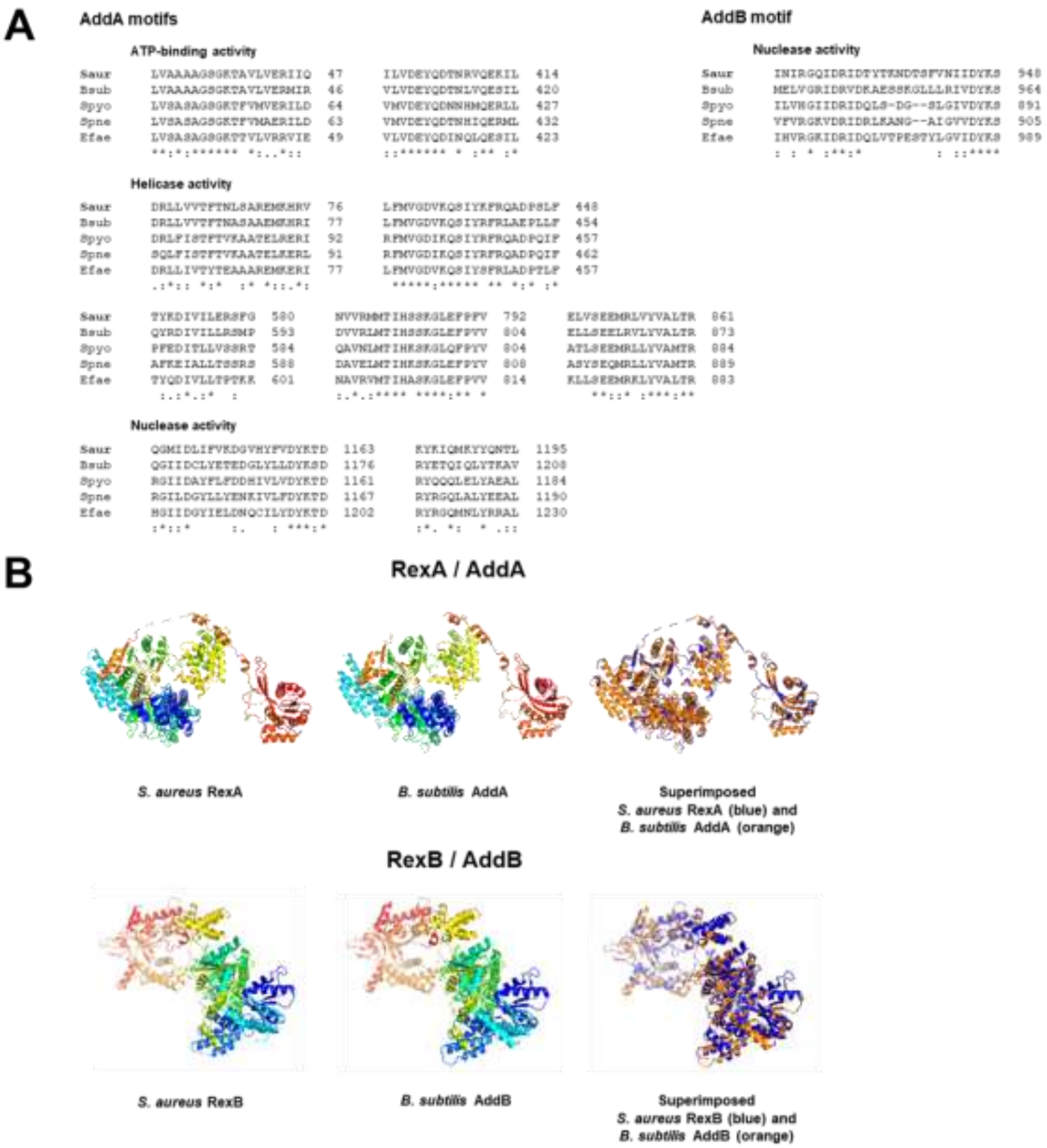


**Figure S4. Structural analysis of *S. aureus* RexAB. (A)** Conserved functional motifs in *S. aureus* RexAB and AddAB homologues. Multiple sequence alignment of conserved motifs from AddA and AddB. AddAB motifs associated with enzyme function were well conserved across the bacterial species examined, with the majority of residues either identical or similar (140 out of 190 in total, 74%). Identical residues are indicated by "*", highly-similar residues by ":" and less similar by ".". Residue positions are indicated on the right-hand side. Alignment was generated by Clustal Omega[1], with protein sequences obtained from the NCBI Protein Database[2]. Saur, *S. aureus*; Bsub, *Bacillus subtilis*; Spyo, *Streptococcus pyogenes*; Spne, *Streptococcus pneumoniae*; Efae, *Enterococcus faecalis*. **(B)** Predicted models of *S. aureus* RexA and RexB superimposed onto individual AddA and AddB subunits of the *B. subtilis* AddAB crystal structure. For *S. aureus* RexA, 1110 residues (91% of the sequence) were modelled with 100% confidence to *B. subtilis* AddA, and for *S. aureus* RexB, 1115 residues (96%) were modelled with 100% confidence to *B. subtilis* AddB. Ribbon representation of *S. aureus* and *B. subtilis* proteins individually and superimposed. Individual protein structures are coloured from blue to red from N- to C-terminals. When superimposed, *S. aureus* proteins are shown in blue and *B. subtilis* proteins in orange. *S. aureus* RexA and RexB predicted 3D models were generated by Phyre2[3]. Protein structures were visualised using PyMOL. PDB 3U4Q was used for the *B. subtilis* AddAB structure[4,5].

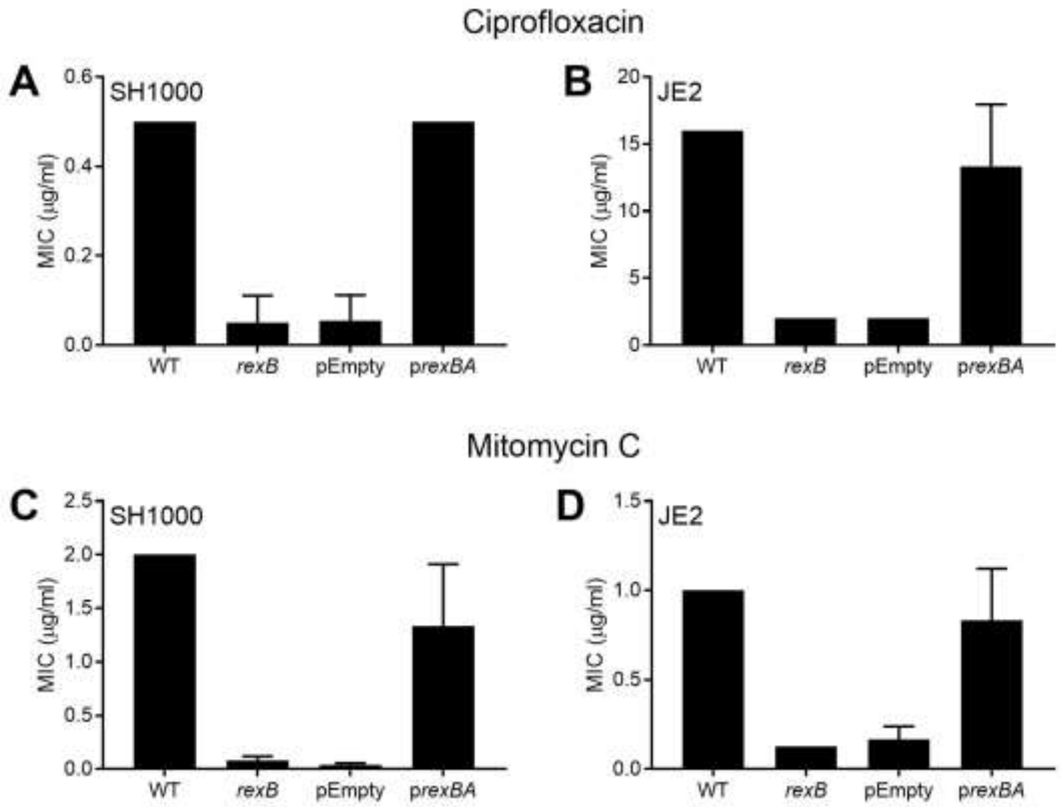


**Figure S5. The *rexBA* mutants are more susceptible than wild type to antibiotics that cause DNA DSBs.** MIC of *S. aureus* WT, *rexB* mutant, empty vector (pEmpty) and complemented mutant (p*rexBA*) in SH1000 and JE2 backgrounds, for ciprofloxacin (A-B) and mitomycin C (C-D). Empty vectors and complemented mutants were supplemented with 100 ng $\mu l^{-1}$ AHT to induce *rexBA* expression (n = 3; median MIC is shown). Error bars represent standard deviation of the mean.

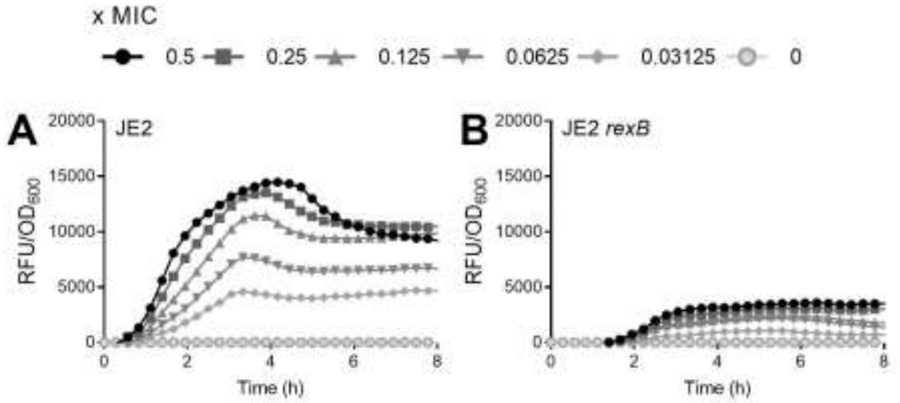


**Figure S6. Mitomycin C causes significantly greater dose-dependent induction of SOS in WT than the *rexB* mutant.** Induction of SOS response of JE2 WT (A) and *rexB* mutant (B) measured by GFP expression upon exposure to varying sub-lethal concentrations of mitomycin C. Concentrations of antibiotic are labelled as multiples of the MIC of the WT strain (x MIC). RFU data are normalised to $OD_{600}$ data to generate *recA* expression relative to cell density (n = 3). Error bars were omitted for clarity.

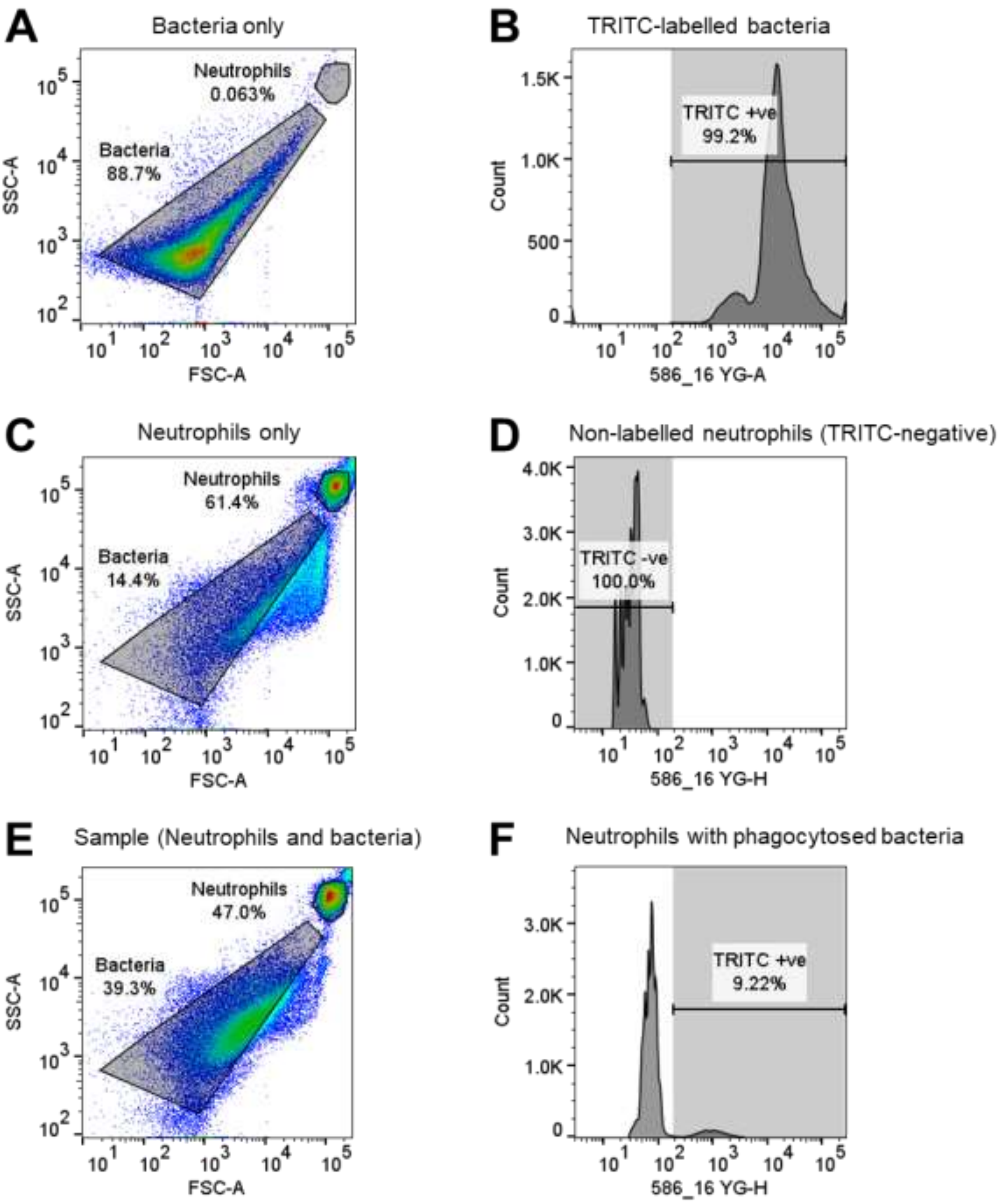


**Figure S7. Gating strategy for flow cytometry data comparing WT and *rexB* mutant *recA-gfp* reporter strains during phagocytosis by neutrophils**. **(A-B)** Bacteria only control gated by forward and side scatter, and subsequently by TRITC labelling. **(C-D)** Neutrophil only control gated by forward and side scatter (C). No TRITC labelling present in the neutrophil sample (D). **(E-F)** Gating strategy used for each sample and time point. Neutrophils and bacteria were gated by forward and side scatter (E). Neutrophils were plotted using 586_16 YG-A for TRITC staining which revealed two distinct populations: neutrophils only (no TRITC detection), and neutrophils containing phagocytosed TRITC bacteria (F). The numbers of phagocytosed bacteria were compared to total number of bacteria to calculate percentage phagocytosis at each time point. To quantify DNA damage, neutrophils containing phagocytosed TRITC bacteria were isolated and the mean GFP value was determined at each time point. Relative GFP detection was used to compare induction of SOS between JE2 WT and *rexB* mutant strains.
